# Helicases DDX5 and DDX17 promote Hepatitis B Virus transcription termination heterogeneity in infected human hepatocytes

**DOI:** 10.1101/2024.01.17.575990

**Authors:** Fleur Chapus, Guillaume Giraud, Pélagie Huchon, Caroline Charre, Chloé Goldsmith, Mélanie Rodà, Maria-Guadalupe Martinez, Judith Fresquet, Audrey Diederichs, Maëlle Locatelli, Hélène Polvèche, Xavier Grand, Caroline Scholtès, Isabelle Chemin, Hector Hernandez Vargas, Michel Rivoire, Cyril F. Bourgeois, Fabien Zoulim, Barbara Testoni

## Abstract

**Background & Aims:** Transcription termination fine tunes gene expression and contributes to specify the function of RNAs in eukaryotic cells. Transcription termination of hepatitis B virus (HBV) is subjected to the recognition of the canonical polyadenylation signal (cPAS) common to all viral transcripts. The regulation of the usage of this cPAS and its impact on viral gene expression and replication is currently unknown.

**Approach & Results:** To unravel the regulation of HBV transcript termination, we implemented a 3’ RACE-PCR assay coupled to single molecule sequencing both in *in vitro* infected hepatocytes and in chronically infected patients. The detection of a previously unidentified transcriptional readthrough indicated that the cPAS was not systematically recognized during HBV replication *in vitro* and *in vivo*. Gene expression downregulation experiments demonstrated a role for the RNA helicases DDX5 and DDX17 in promoting viral transcriptional readthrough, which was, in turn, associated to HBV RNA destabilization and decreased HBx protein expression. RNA and chromatin immunoprecipitation, together with mutation of cPAS sequence suggested a direct role of DDX5 and DDX17 in functionally linking cPAS recognition to transcriptional readthrough, HBV RNA stability and replication.

**Conclusions:** Our findings identify DDX5 and DDX17 as crucial determinants for HBV transcriptional fidelity and as host restriction factors for HBV replication.

## INTRODUCTION

Despite treatments efficient in maintaining the infection under control, Hepatitis B virus (HBV) cannot be fully eradicated from infected hepatocytes (1). Viral persistence is due to the so-called covalently closed circular DNA (cccDNA) that acts as a viral minichromosome in the nucleus of infected hepatocytes. cccDNA is the unique template for the transcription of the six main viral mRNAs by the host RNA polymerase II (RNAP II), including the pregenomic RNA (pgRNA). The latter is then retro-transcribed and encapsidated to generate new infectious particles. Thus, silencing transcriptional activity of cccDNA is considered as a promising therapeutic approach to achieve a functional cure of chronic HBV infection (2).

Similarly to host transcripts, HBV RNAs are subjected to a series of co-transcriptional processing events that not only contribute to their maturation, but also regulate HBV replication and/or contribute to HBV-induced liver diseases (3). Co-transcriptional HBV RNA polyadenylation is initiated by the recognition of a polyadenylation signal (cPAS) common to all HBV transcripts as a termination signal for the RNAP II. Its sequence, UAUAAA, is derived from the consensus mammalian PAS sequence AAUAAA (4). Furthermore, *in vitro* assays showed that both 5’ and 3’ PAS-surrounding sequences are critical for its proper recognition and thus for a proper 3’ end processing of HBV transcripts (4, 5). However, the molecular mechanisms governing proper transcription termination and how it influences HBV replication remain open questions.

RNA helicases play fundamental roles in the regulation of gene expression, including transcription termination and 3’ end mRNA processing (6). Members of this protein family appeared to act as enhancer or repressor of HBV replication (7). Among these proteins, the highly redundant ATP-dependent RNA helicases DDX5 and DDX17 were shown to couple transcription and co-transcriptional mRNA processing to fine tune gene expression during differentiation processes (6, 8). In particular, DDX5 and DDX17 act as regulators of transcription termination and 3’ end mRNA processing in several cellular models (9, 10). DDX5 was also identified as a HBV restriction factor contributing to silence cccDNA transcriptional activity (11), while the role of DDX17 in HBV lifecycle still remains controversial (12, 13). Noteworthy, these previous studies were performed in the transformed hepatoma cell lines HepG2-NTCP.

Here, in addition to HepG2-NTCP cells, we used non transformed primary human hepatocytes (PHH), as well as patients’ liver tissue samples, and combined 3’ Rapid amplification of cDNA ends (RACE)-PCR and single molecule sequencing using the Oxford Nanopore Technology (ONT) MinION to accurately map the 3’ extremity of HBV transcripts *in vitro* and *in vivo*. We demonstrated that, while most of HBV RNAs end 14 nucleotides downstream of the HBV cPAS, a proportion of viral transcripts display a transcriptional readthrough spanning around 700 nucleotides. Inhibition of this readthrough following DDX5/17 silencing identified DDX5 and DDX17 as critical regulators of HBV cPAS recognition. Furthermore, we demonstrated that this transcriptional readthrough promoted by DDX5 and DDX17 was associated with HBV RNAs destabilization. Accordingly, we provide evidence that DDX5/17 knockdown and, consequently, the preferential usage of the HBV cPAS were associated with enhanced HBV replication, establishing a functional link between transcriptional termination and proper viral replication.

## EXPERIMENTAL PROCEDURES

### Ethics statement

Primary human hepatocytes (PHH) were isolated from surgical liver resections after informed consent of patients (IRB agreements #DC-2008-99 and DC-2008-101).

The protocol for the use of liver samples from two CHB patients was approved by the competent Institutional Ethics Committee (CPP Sud est IV 11/040, authorization number DC-2008-235). Written informed consent was obtained from all patients and/or their legal guardians to underwent liver biopsy.

### Additional materials and methods

All experimental procedures and materials are available on supplementary information.

### Statistical analysis

Data are expressed as mean ± standard error of the mean (SEM). Mann-Whitney or Kruskal-Wallis (multiple comparison) tests were used when appropriate (p<0.05).

## RESULTS

### Mapping of HBV transcripts 3’ extremity reveals differential recognition of polyadenylation signal

We used HepG2-NTCP cells and primary human hepatocytes (PHHs), two validated cellular models to study HBV replication, to characterize the 3’ extremity of viral transcripts 8 days post-infection, when viral replication is well established (14–16). We set-up an HBV-specific 3’ RACE approach coupled to PCR that, given the overlapping nature of HBV RNAs, allows the amplification of the 3’ end of all viral transcripts (**Figure 1a**). Agarose gel electrophoresis revealed a smeared pattern in both HBV-infected HepG2-NTCP cells and PHHs, with recognizable discrete bands between 400 bp and 700 bp in size (**Figure S1a**). Nevertheless, longer amplicons reaching up to 1200 bp were also observed in both cellular models (**Figure S1a**), highlighting an unexpected heterogeneity in the length of the 3’ region of HBV transcripts.

**Figure 1:**
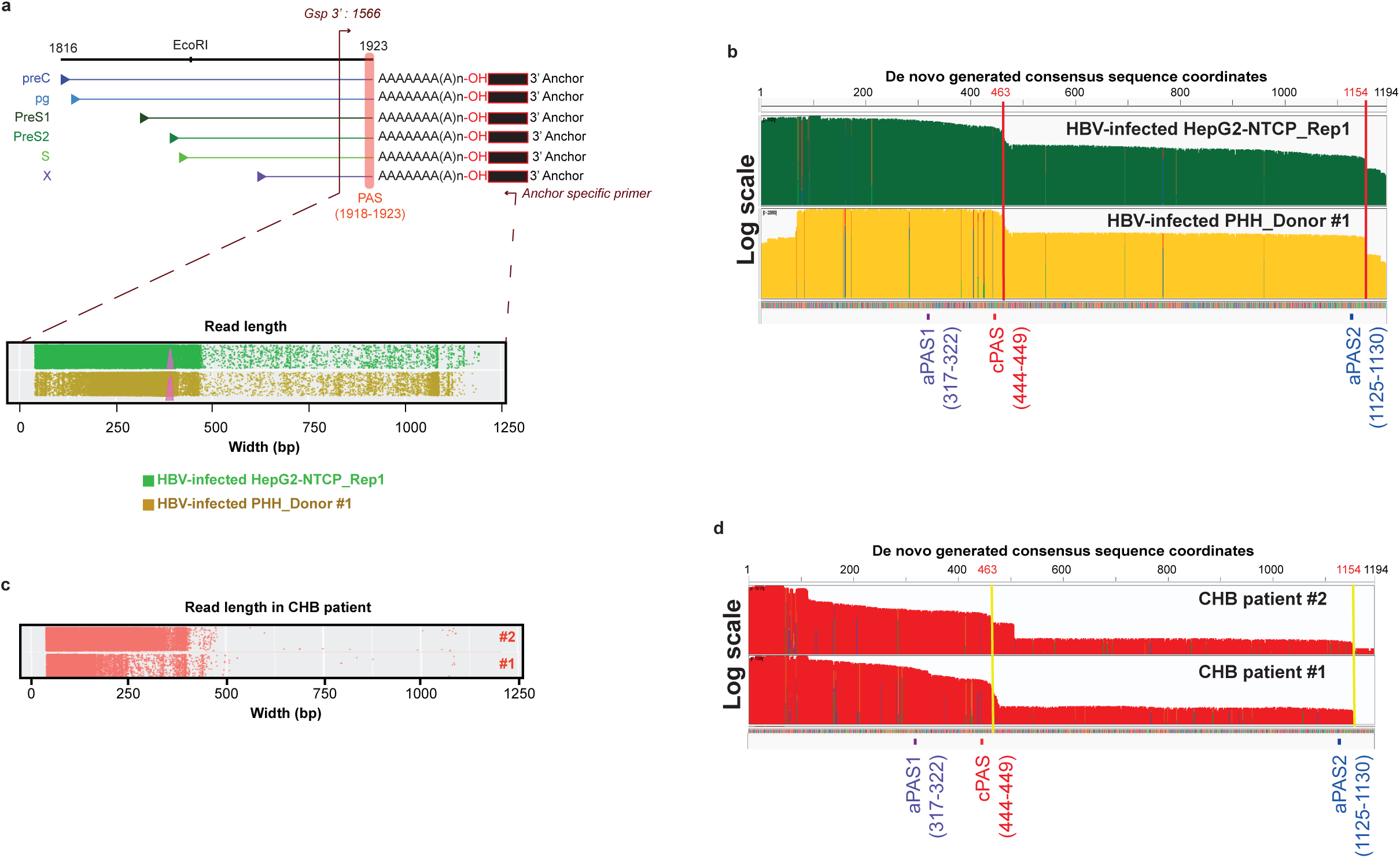
HBV transcripts end 14 nucleotides downstream of the cPAS and display transcriptional read-through. RNAs extracted from HepG2-NTCP cells and primary human hepatocytes (PHH) infected with HBV at MOI 250 for 8 days (a-b) and from two CHB patients’ liver resections (c-d) were subjected to HBV-specific 3’RACE-PCR experiments followed by MinION single molecule sequencing. **a, c** Graphical representation of the reads length obtained from the MinION single molecule sequencing of 3’RACE-PCR amplicons. Each dot corresponds to a single read. **b**, **d** Integrated Genome Viewer (IGV) view of the Nanopore sequencing reads aligned to a *de novo* assembly generated HBV consensus sequence. At the bottom, the position of poly A sites (PAS) predicted by the Poly (A) Signal Miner prediction tool is indicated. The PAS indicated in red (common PAS, cPAS) corresponds to the previously identified PAS known to be shared by every HBV transcripts (4). The vertical lines (red or yellow) correspond to the end of HBV transcripts. The scale used is a logarithmic scale. aPAS, alternative PAS.

Sequencing profiles obtained by single molecule ONT MinION analysis of 3’ RACE amplicons confirmed the previous observation. Indeed, a significant proportion of reads reaching up to ∼ 1100 bp were observed, besides those with a size of around 450 bp. The latter corresponds to the expected amplicon size if the cPAS was systematically used for viral transcript termination (**Figure 1a lower panel, S1b**).

To better characterize these longer reads, a consensus sequence of 1194 bp was generated by *de novo* assembly of the totality of the sequencing reads. The generated consensus was consistent across biological replicates and showed 98 % identity with HBV genotype D genome by BLAST analysis (**Figure S1c**). We then aligned every single sequencing read to this reference and visualized them using Integrated Genomics Viewer (IGV) (17) (**Figure 1b**). Similarly to eukaryotic cell transcripts, which are cleaved between 10 and 30 nucleotides downstream of the PAS (18), we located the main HBV RNAs termination site 14 nucleotides downstream of the previously described cPAS in both HepG2-NTCP cells and in PHHs (**Figure 1b, S1d**) (4). These data are in agreement with the main use of this cPAS as a termination signal for RNAP II and concordant with previous *in vitro* work (4). Sequencing analysis confirmed that the longer reads shown in **Figure 1b** correspond indeed to single HBV transcripts extending up to 705 bp downstream of the cPAS in both cellular models (**Figure 1b, S1d)**, indicating a transcriptional readthrough. We looked for the consensus PAS sequence, NNTANA, obtained from the 12 mostly used PAS in human genome, to determine whether potential alternative PAS (aPAS) could be used to terminate the transcription of these longer transcripts (19) (**Figure S1e**). Besides the known cPAS (at the position 1923 of the HBV genome) and the upstream aPAS1 (position 1800) (20), several other consensus PAS sequences were bioinformatically identified in the transcriptional readthrough sequence, including the aPAS2 located 19 nucleotides upstream of the major end of the longer transcripts (at the position 2603 of the HBV genome) suggesting that aPAS2 could potentially be recognized as a termination signal (**Figure 1b, S1d,e**).

Similar analyses performed in liver biopsies derived from two CHB patients confirmed the results obtained in cell models, with a major peak of reads of 450 bp corresponding to HBV transcripts ending 14 nucleotides downstream of the HBV cPAS (**Figure 1c-d, S1f**) and longer reads of different sizes going up to 705 bp downstream of the HBV cPAS (**Figure 1c-d**).

Altogether, our data showed that HBV transcripts end 14 nucleotides downstream of the previously described HBV cPAS, but also pointed to the existence of a transcriptional readthrough terminating downstream of a putative aPAS, demonstrating that the viral cPAS is not systematically recognized as the unique termination signal for RNAP II.

### DDX5 and DDX17 favour the HBV transcriptional readthrough

The uncovering of HBV transcriptional readthrough led us to investigate the mechanisms regulating HBV cPAS recognition. Transcription termination and 3’ end mRNA processing are tightly linked and regulated at both the chromatin and the RNA level (21). The ATP-dependent RNA helicases DDX5 and DDX17 play major roles in coupling transcription and co-transcriptional mRNA processing and were identified as part of protein complexes containing termination factors (6, 8–10).

To investigate if DDX5 and DDX17 regulate the recognition of the HBV cPAS, we concomitantly repressed DDX5 and DDX17 expression by RNAi in HBV-infected HepG2- NTCP cells 8 dpi (**Figure S2a-b**). 3’RACE-PCR followed by agarose gel electropohoresis revealed several bands in the control RNAi condition (**Figure S2c**), while in DDX5/17-silenced HBV-infected HepG2-NTCP cells, a major band of around 500 bp was detected, suggesting a loss of heterogeneity at HBV 3’ transcripts region when DDX5 and DDX17 were downregulated (**Figure S2c**). ONT MinION single molecule sequencing of 3’RACE amplicons revealed that HBV-infected HepG2-NTCP cells transfected with non-targeting siRNA displayed the same pattern as non-transfected cells, with reads reaching up to around 1200 bp (**Figure 2a, S2d**). Interestingly, these reads appeared to be significantly less frequent in DDX5/17-depleted HepG2-NTCP cells compared to control cells (**Figure 2a-b, S2d**-**e**). We then aligned every read to the previously *de novo* generated HBV reference genome and confirmed that the depletion of DDX5 and DDX17 decreased the frequency of transcriptional readthrough in favour of a termination 14 nt downstream of the cPAS (**Figure 2c, S2f**).

**Figure 2:**
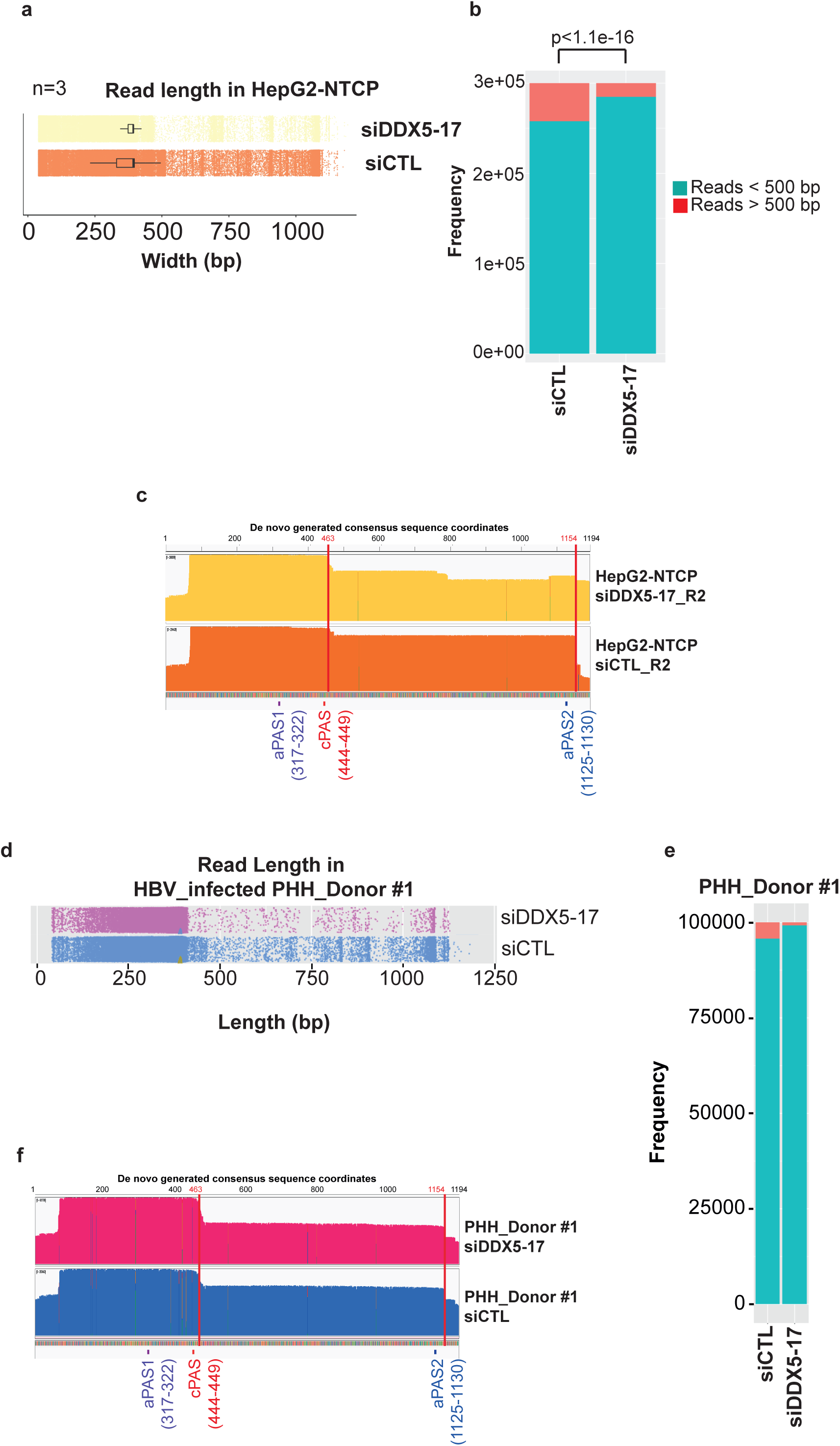
DDX5 and DDX17 favour the transcriptional readthrough. HepG2-NTCP cells and PHH were infected with HBV at MOI 250 for 8 days. **a, d** Graphical representation of the reads length obtained from 3 independent MinION sequencing experiments of the 3’RACE-PCR amplicons obtained from HBV-infected HepG2- NTCP cells (**a**) or PHH (**d**) transfected with control siRNA (siCTL) or DDX5/17 targeted siRNAs (siDDX5-17). Each dot corresponds to a single read. **b, e** Barplot indicating the frequency of the reads longer (in red) or shorter (in blue) than 500 bp obtained from the single molecule sequencing of the 3’RACE amplicons showed in **Figure S2c and S2i,** respectively. Fisher test comparing the proportion of long to short reads, α threshold=0.05. **c, f** IGV view of the MinION reads aligned to the *de novo* assembly generated HBV consensus reference genome and obtained from the HBV-infected HepG2-NTCP cells (**c**) or PHH (**f**) transfected with control siRNA (siCTL) or with DDX5/17-targeted siRNA.

Similar experiments were performed in HBV-infected PHH derived from two different donors (**Figure 2d-f and S2g-l**). As observed in the HepG2-NTCP model, HBV transcripts displayed a reproducibly less frequent transcriptional readthrough in DDX5/17-depleted cells compared to control PHH, (**Figure 2d-f and S2g-l**), confirming that DDX5 and DDX17 also prevented the recognition of the HBV cPAS in infected primary cells.

Altogether, these data showed that DDX5 and DDX17 are involved in HBV mRNA 3’ end processing by preventing the recognition of the cPAS and promoting HBV transcriptional readthrough.

### DDX5 and DDX17 bind cccDNA and HBV RNAs

To determine whether DDX5 and DDX17 act directly to promote HBV transcriptional readthrough, we first analysed the transcriptome of HBV-infected HepG2-NTCP cells depleted or not for both RNA helicases by RNA-seq experiments. DEseq2 gene expression analysis identified 1585 up-regulated and 2088 down-regulated genes after the repression of DDX5 and DDX17 (log_2_foldchange > |1|). Gene ontology analyses performed using GenePattern website (https://www.genepattern.org) did not identify terms related to 3’ end mRNA processing or transcription termination suggesting that DDX5 and DDX17 do not regulate genes involved in this process in the experimental conditions analysed (**Figure 3a**).

**Figure 3:**
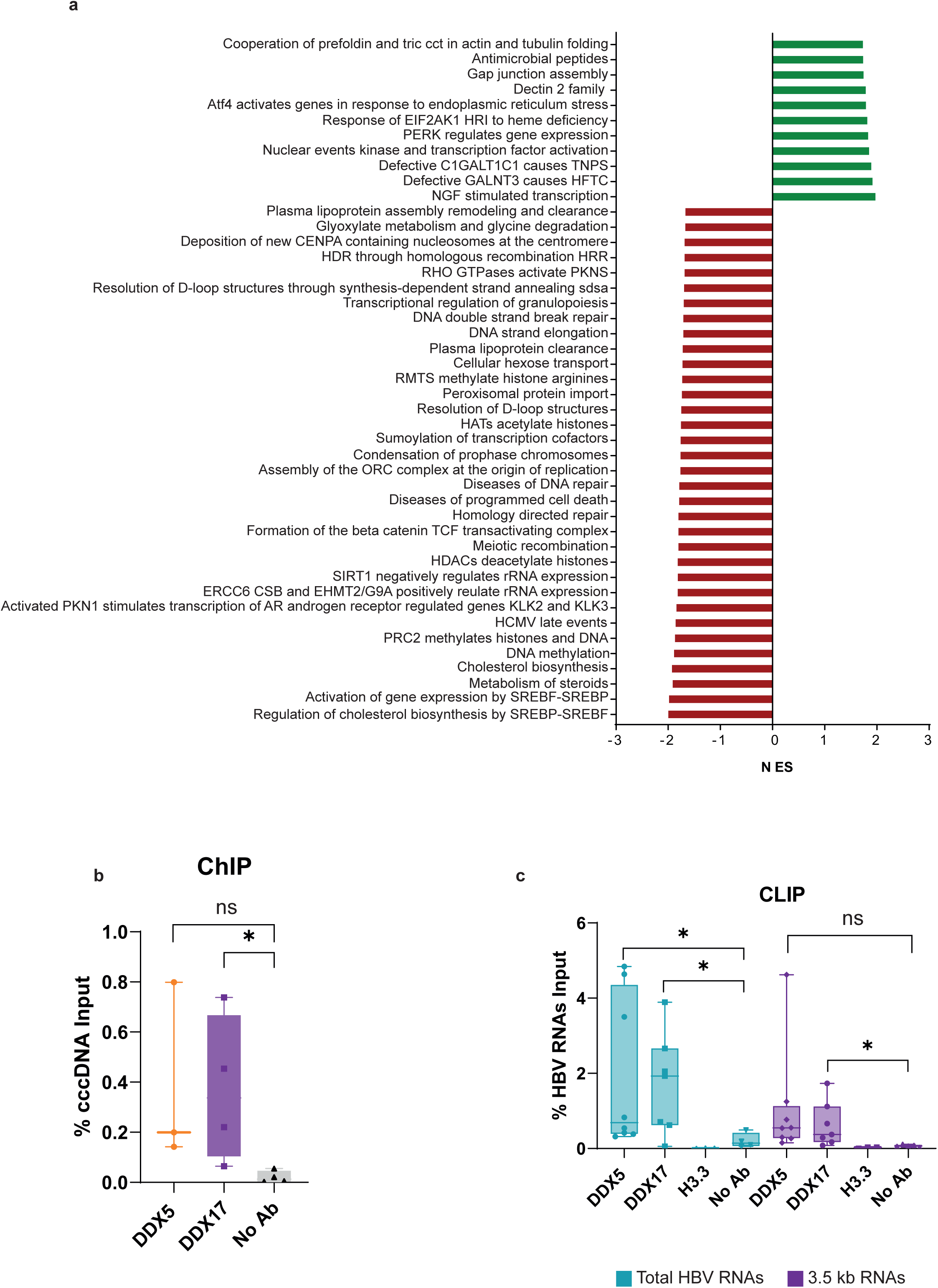
Direct role of DDX5 and DDX17 in the regulation of HBV transcription termination. **a** Signalling pathways enriched from the list of up-regulated (green bars) and down-regulated (red bars) genes after DDX5 and DDX17 repression in HBV-infected HepG2-NTCP identified by RNA-Seq followed by gene ontology analyses using GenePattern. **b** ChIP experiments were performed from crosslinked HBV-infected HepG2-NTCP cells using an anti-DDX5 (orange), an anti-DDX17 (purple) or without antibody (NoAb, black) followed by qPCR experiments to amplify precipitated cccDNA. The graph represents the percentage of cccDNA present in the Input material. Each dot represents one biological replicate. Boxplots represent the average of the three (DDX5) or four (DDX17 and NoAb) independent biological replicates and the error bars represent the standard error to the mean (s.e.m.). *: Mann-Whitney p-value <0.05. **c** CLIP experiments were performed from UV-crosslinked HBV-infected HepG2-NTCP cells using an anti-DDX5, an anti-DDX17 or an anti-H3.3 antibody or without antibody (No Ab) followed by RT-qPCR experiments with assays detecting all HBV transcripts (total HBV RNAs, green) or 3.5 kb RNAs (purple). The graph represents the percentage of HBV RNAs present in the Input material. Each dot represents one biological replicate. Boxplots represent the average of two (H3.3) or six (DDX5, DDX17 and NoAb) independent biological replicates and the error bars represent the standard error of the mean (s.e.m.). *: Mann-Whitney p-value <0.05.

In parallel, we investigated the ability of both RNA helicases to bind cccDNA and HBV RNAs. Chromatin immunoprecipitation (ChIP)-qPCR (**Figure 3b**) and CLIP (UV-crosslinking immunoprecipitation) (RT-)qPCR (**Figure 3c**) were carried out in HBV-infected HepG2-NTCP cells. cccDNA and HBV RNAs showed a specific enrichment compared to negative controls (NoAb for ChIP and H3.3 and NoAb for CLIP) when antibodies against either DDX5 or DDX17 were used for Immunoprecipitation (IP) (**Figure 3b-c**). These data demonstrate that DDX5 and DDX17 are recruited to HBV cccDNA and RNAs and suggest that they could play a role in the recognition of cPAS and, thus, in the regulation of HBV 3’ end mRNA processing.

### Lengthening of the 3’UTR destabilizes HBx transcripts

The transcriptional readthrough of HBV transcripts results in the lengthening of their 3’ untranslated region (3’ UTR). 3’ UTR is a critical regulatory region for RNA metabolism and the site of recruitment of regulators such as RNA binding proteins or miRNAs, known to dictate RNA stability, export and/or translation (22).

To unveil the impact of the transcriptional readthrough on HBV RNA metabolism, we focused on its effect on HBx transcript, which codes for the HBx protein, indispensable for HBV infection (23). To mimic the lack of cPAS recognition observed in DDX5/17 depleted condition, we engineered expression vectors (pcDNA6) bearing the HBx coding sequence fused at its 5’ end to a V5 tag coding sequence and at its 3’ end to the 3’ UTR region encoded by the transcriptional readthrough carrying a WT (cPASwt) or mutated cPAS (TATAAA to CGCGGG, cPASmut) (**Figure 4a**).

**Figure 4:**
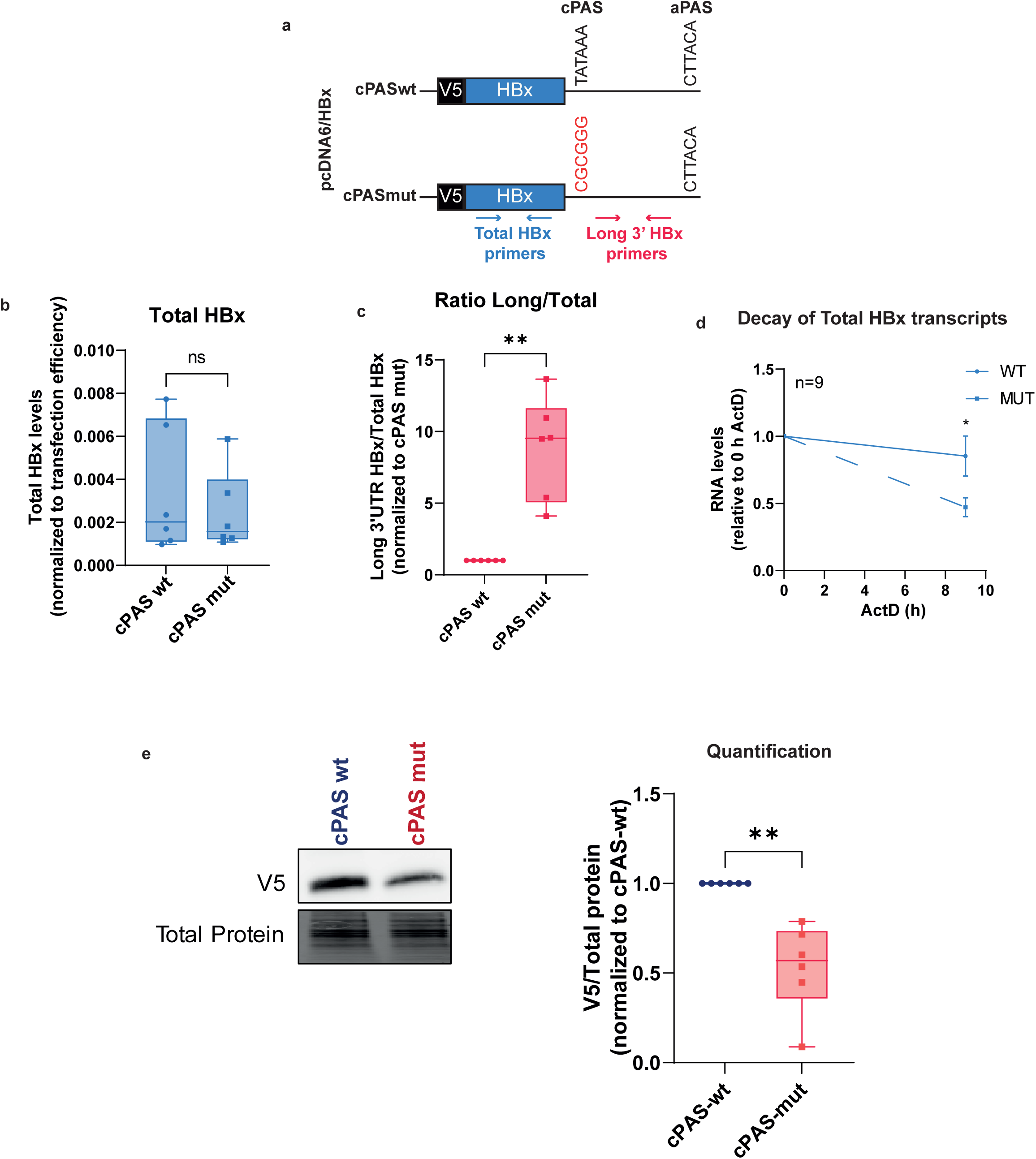
HBV transcriptional readthrough destabilizes HBx transcripts. HepG2-NTCP cells were transfected with either the cPASwt or the cPASmut constructs (**a**). Two days after, RNAs and proteins were collected and analysed by RT-qPCR and Western blot, respectively. **b, c** Quantification of HBx transcripts with assays recognizing all forms (total, **b**) and only long (**c**) HBx RNAs by RT-qPCR. Levels of total HBx transcripts were normalized to the levels of *GAPDH* mRNAs and of transfected pcDNA6 plasmids quantified by qPCR (**Figure S3b**). Levels of long HBx transcripts were normalized to the levels of total HBx transcripts. Each dot represents one biological replicate. Boxplots represent the first quartile, the median (horizontal line) and the third quartile of six independent biological replicates. Error bars represent the minimal and maximal values. **d** cPASwt (full lines) and cPASmut (dashed lines)-transfected HepG2-NTCP cells were treated with 20 µg/mL ActD for 9 h. RNAs were extracted at 0 and 9 hours post-treatment and subjected to RT-qPCR to amplify total HBx transcripts. Each time point represents the average <u>±</u> standard error of the mean (SEM) of nine independent biological replicates. **e** Quantification of V5 protein levels by Western blot analyses using an anti-V5 antibody. Total proteins blots used as loading controls were obtained after activation of the stain-free gels. Boxplots represent the densitometry quantification of the V5 signal normalized to the corresponding signal for total proteins and expressed as fold change respect to cPASwt transfected condition. ns: Mann-Whitney p-value > 0.05; *: Mann-Whitney p-value < 0.05; **: Mann-Whitney p-value < 0.01; ****: Mann-Whitney p-value <0.0001.

After transfection of HepG2-NTCP cells (**Figure S3a**), the levels of total HBx transcripts and HBx transcripts bearing a long 3’UTR (long HBx) were quantified by RT-qPCR experiments. While the levels of total HBx transcripts produced by the cPASmut constructs did not appear to be significantly different from the cPASwt (**Figure 4b**), the cPASmut construct produced a significantly increased proportion of long HBx transcripts than the cPASwt construct (**Figure 4c**). Strikingly, long HBx transcripts produced by the cPASwt construct could still be detected, further confirming that the cPAS is not systematically recognized as a termination signal for RNAP II in this construct, similarly to HBV genome after infection (**Figure 1, 4c**).

We then tested the impact of the long 3’UTR on HBx transcripts stability. We treated cPASwt- and cPASmut-transfected HepG2-NTCP cells with Actinomycin D (ActD), a transcription inhibitor, for 9 hours and quantified the remaining levels of total HBx transcripts by RT-qPCR. We observed a significantly faster decay of total HBx transcripts produced by the cPASmut constructs compared to those produced by the cPASwt construct (**Figure 4d**). Since we observe a significant increase of RNAs presenting a readthrough in the cPASmut condition, these data strongly suggest that the presence of a longer 3’UTR destabilizes HBx transcripts.

Finally, we compared HBx protein levels produced by cPASwt- and cPASmut- transfected HepG2-NTCP cells. Western blot analyses using an anti-V5 antibody revealed that the cPASmut construct produced significantly less HBx protein than the cPASwt construct (**Figure 4e**), strongly associating the presence of longer 3’UTR to HBx RNA destabilization and lower HBx protein levels.

### DDX5 and DDX17 repress HBx by promoting transcriptional readthrough

To establish a functional link between DDX5-17 expression and the transcriptional readthrough, we tested the effect of DDX5 and DDX17 up- or down-regulation on the HBx RNA and protein levels in cPASwt and cPASmut-transfected HepG2-NTCP cells (**Figure S4a-b**). Silencing of DDX5 and DDX17 increased the global level of total HBx transcripts produced by the two constructs, in accordance with a previously described role for DDX5 as HBV transcriptional repressor(11) (**Figure 5a**). However, the proportion of long HBx transcripts was decreased only when the cPASwt construct was transfected, whereas it was not affected by DDX5-17 silencing in the cPASmut condition (**Figure 5b**). HBx protein levels in the siCTL cPASmut condition were lower than in the corresponding siDDX5-17 (**Figure 5c**), thus confirming the results obtained previously (**Figure 4e**). Nevertheless, DDX5-17 silencing induced an increase of HBx protein produced from both constructs, thus reflecting the mixed effect of both the increase in total HBx transcripts and the decrease of the transcriptional readthrough (**Figure 5c**). Overexpression of DDX5 and DDX17 either individually (**Figure S4c-e**) or in combination (**Figure 5d-f**) did not affect the levels of total HBx transcripts (**Figure 5d**), but significantly increased the proportion of long HBx RNAs produced from the cPASwt construct but not from the cPASmut construct (**Figure 5e**). This imbalanced proportion of longer HBx transcripts in the cPASmut condition was associated to decreased levels of HBx protein (**Figure 5f**).

**Figure 5:**
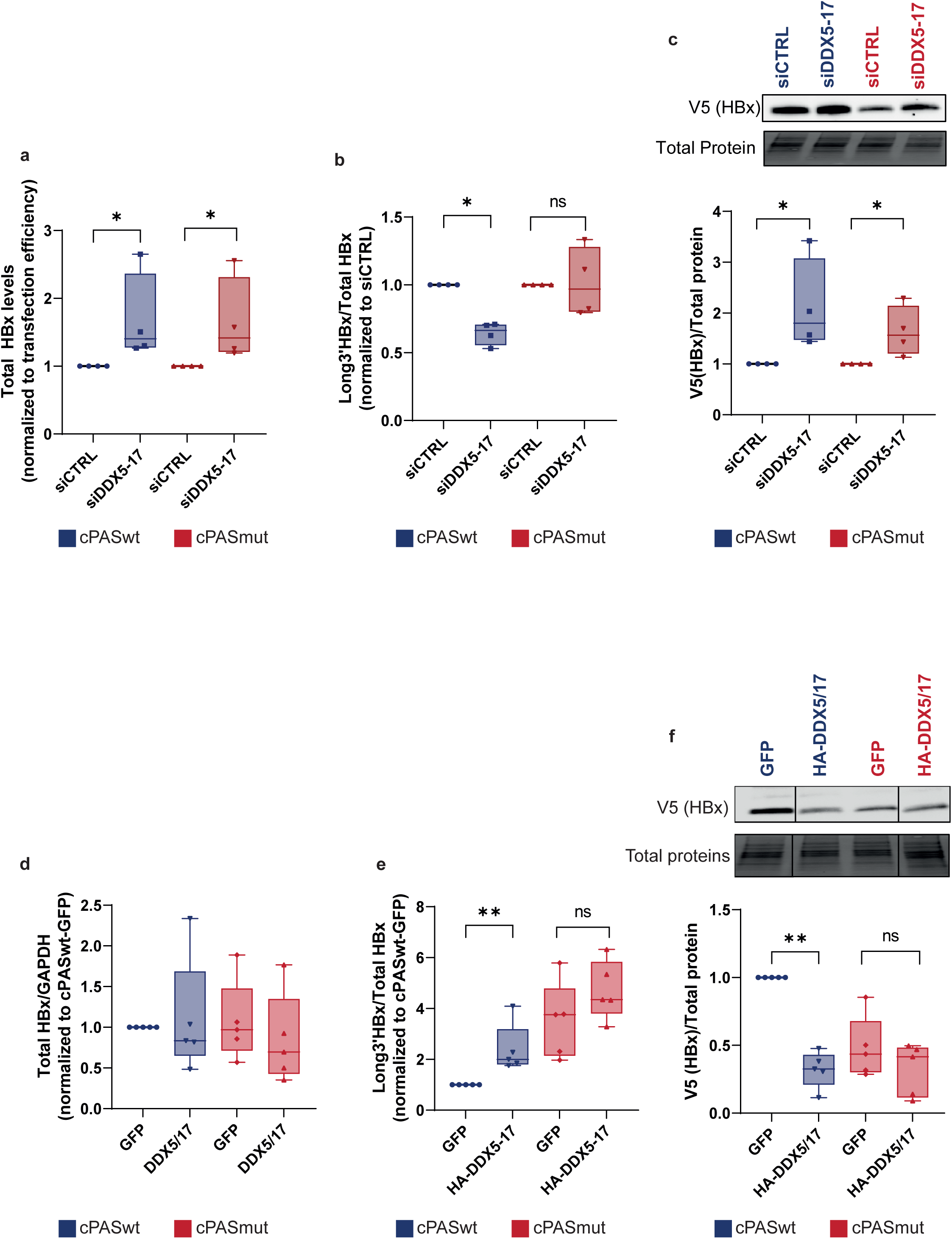
DDX5 and DDX17 regulate HBx expression by promoting transcriptional readthrough. **a, b, c** HepG2-NTCP cells were transfected with control siRNAs (siCTRL) or siRNAs directed against *DDX5* and *DDX17* mRNAs one day prior to their transfection with either cPASwt or cPASmut constructs. Two days after the first transfection, RNAs and proteins were extracted and subjected to RT-qPCR (**a, b**) and Western blot analyses (**c**) to quantify long HBx transcripts (**a**), total HBx transcripts (**b**) and HBx protein levels (**c**). **d, e, f** HepG2-NTCP cells were transfected with a GFP or HA-DDX5 and HA-DDX17 expression plasmids one day prior to their transfection with either cPASwt or cPASmut constructs. Two days after the first transfection, RNAs and proteins were extracted and subjected to RT-qPCR (**d, e**) and Western blot analyses (**f**) to quantify total HBx transcripts (**d**), long HBx transcripts (**e**) and HBx protein levels (**f**). **a, e** Long HBx transcripts levels were normalized to total HBx transcripts levels and the siCTRL-cPASwt (**a**) or the GFP-cPASwt (**e**) conditions. **b, d** Total HBx transcripts levels were normalized to *GAPDH* levels and the siCTRL-cPASwt (**b**) or the GFP-cPASwt (**d**) conditions. **c, f** V5 (HBx) signal was quantified and normalized to the corresponding total proteins load and siCTRL-cPASwt (**c**) or GFP-cPASwt (**f**) conditions. **a-e** Each dot represents one replicate. Boxplots represent the first quartile, the median (horizontal line) and the third quartile of four or five independent biological replicates. Error bars represent the minimal and maximal values. ns: Mann-Whitney p-value > 0.05; *: Mann-Whitney p-value < 0.05. **: Mann-Whitney p-value < 0.01.

Altogether, these data strongly suggest that DDX5 and DDX17 repress HBx protein expression at least by promoting the transcriptional readthrough through a mechanism dependent on the recognition of cPAS sequence.

### DDX5 and DDX17 silencing stabilizes HBV RNAs in HBV-infected HepG2-NTCP cells and promotes viral replication

Then, we investigated if DDX5 and DDX17 were involved in the regulation of the stability of HBV RNAs produced in HBV-infected HepG2-NTCP cells. Indeed, treatment of HBV-infected HepG2-NTCP cells with ActD for 9 hours revealed a significantly slower decay of total HBV RNAs after knockdown of DDX5 and DDX17 compared to control cells (**Figure 6a, left panel**). Surprisingly, the slower decay was less pronounced for 3.5 kb RNAs and appeared not to be statistically significant (**Figure 6a, central panel)**. *GUSB* transcript half-life was not impacted by DDX5 and DDX17 silencing (**Figure 6a, right panel**), indicating a specific effect of DDX5 and DDX17 silencing on HBV RNAs. Accordingly, knockdown of DDX5 and DDX17 induced a significant increase of total HBV RNAs levels (**Figure 6b**). The levels of 3.5 kb RNAs were, instead, only slightly increased (**Figure 6b**), prompting us to investigate if this was due to a higher rate of retrotranscription associated to the degradation of pgRNA mediated by the RNAseH activity of the viral polymerase. To this aim, we first quantified by qPCR the levels of cytoplasmic encapsidated HBV DNA after nucleocapsid isolation and showed that DDX5 and DDX17 repression was associated with a significant increase of encapsidated HBV DNA levels in the cytoplasm of infected cells (**Figure 7a**). In the same samples, Native Agarose Gel Electrophoresis (NAGE) coupled to western blotting analysis with anti-capsid (HBc) and envelope (HBs) HBV proteins revealed that the level of both cytoplasmic non-enveloped and enveloped capsids signal was also increased, thus showing enhanced virion formation and viral replication upon DDX5/17 downregulation (**Figure 7b**). This was accompanied by stable levels of cccDNA detected both by qPCR and Southern blotting (**Figure 7c and S5**).

**Figure 6:**
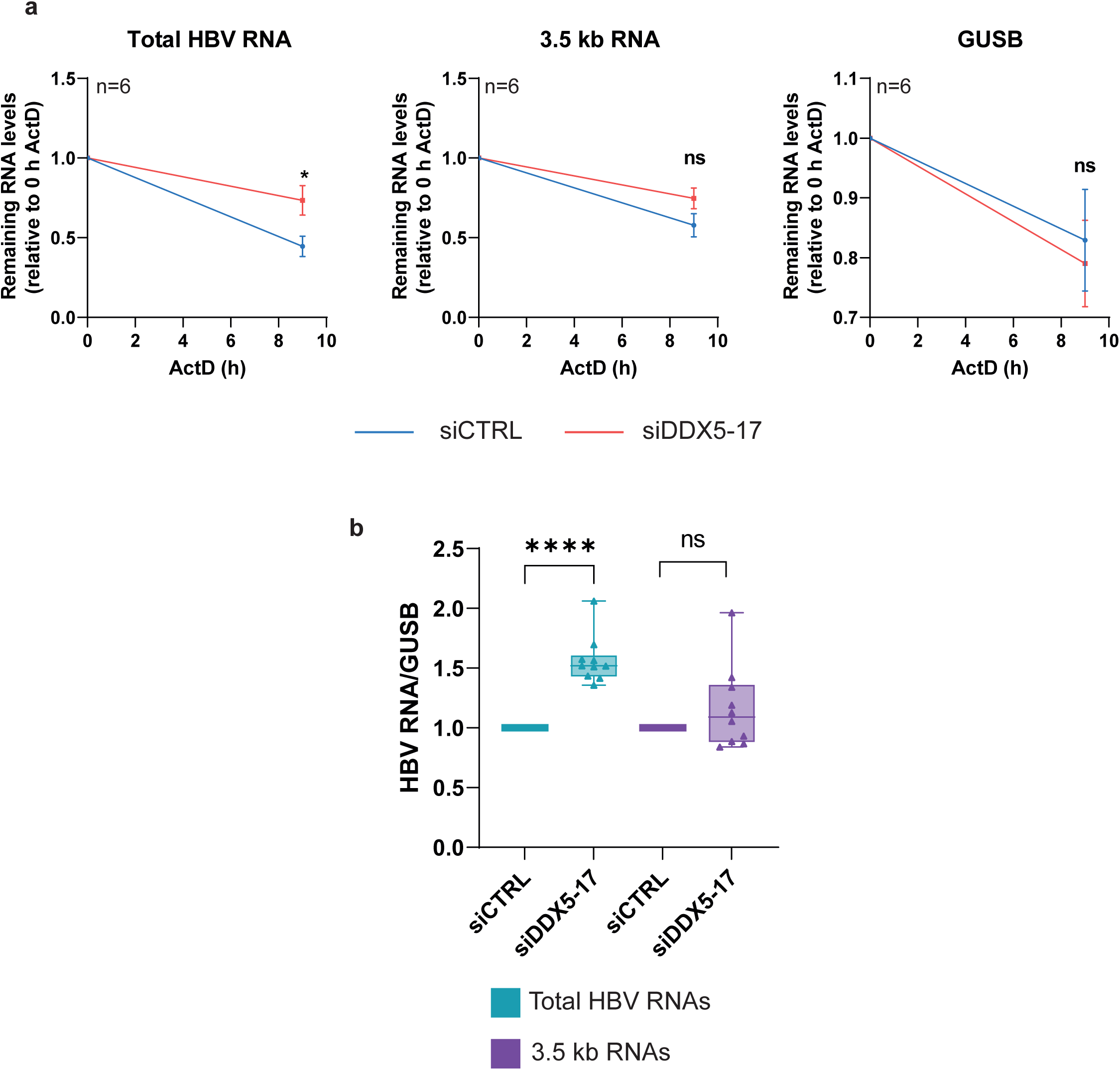
DDX5 and DDX17 repress and destabilize HBV RNAs. HepG2-NTCP cells were infected at MOI 250 and transfected at 4 and 6 days post-infection with control siRNA (siCTRL) or siRNAs directed against *DDX5* and *DDX17* mRNAs (siDDX5-17). Eight days post-infection, RNAs were extracted and subjected to RT-qPCR analyses. **a** 20 µg/mL ActD were added or not 9 h prior to RNA extraction. The graphs represent the quantification of remaining levels of total HBV RNAs (left panel), 3.5 kb RNAs (center panel) or *GUSB* (right panel). Each time point represents the average of six independent biological replicates ± SEM. **b** Quantification of total HBV RNAs (green) or 3.5 kb RNAs (purple). Levels of HBV RNAs were normalized to *GUSB* levels and expressed as fold change of the siCTRL condition. Boxplots represent the first quartile, the median (horizontal line) and the third quartile of ten independent biological replicates. Error bars represent the minimal and maximal values. ns: Mann-Whitney p-value > 0.05; *: Mann-Whitney p-value < 0.05; **: Mann-Whitney p-value < 0.01; ****: Mann-Whitney p-value <0.0001.

**Figure 7:**
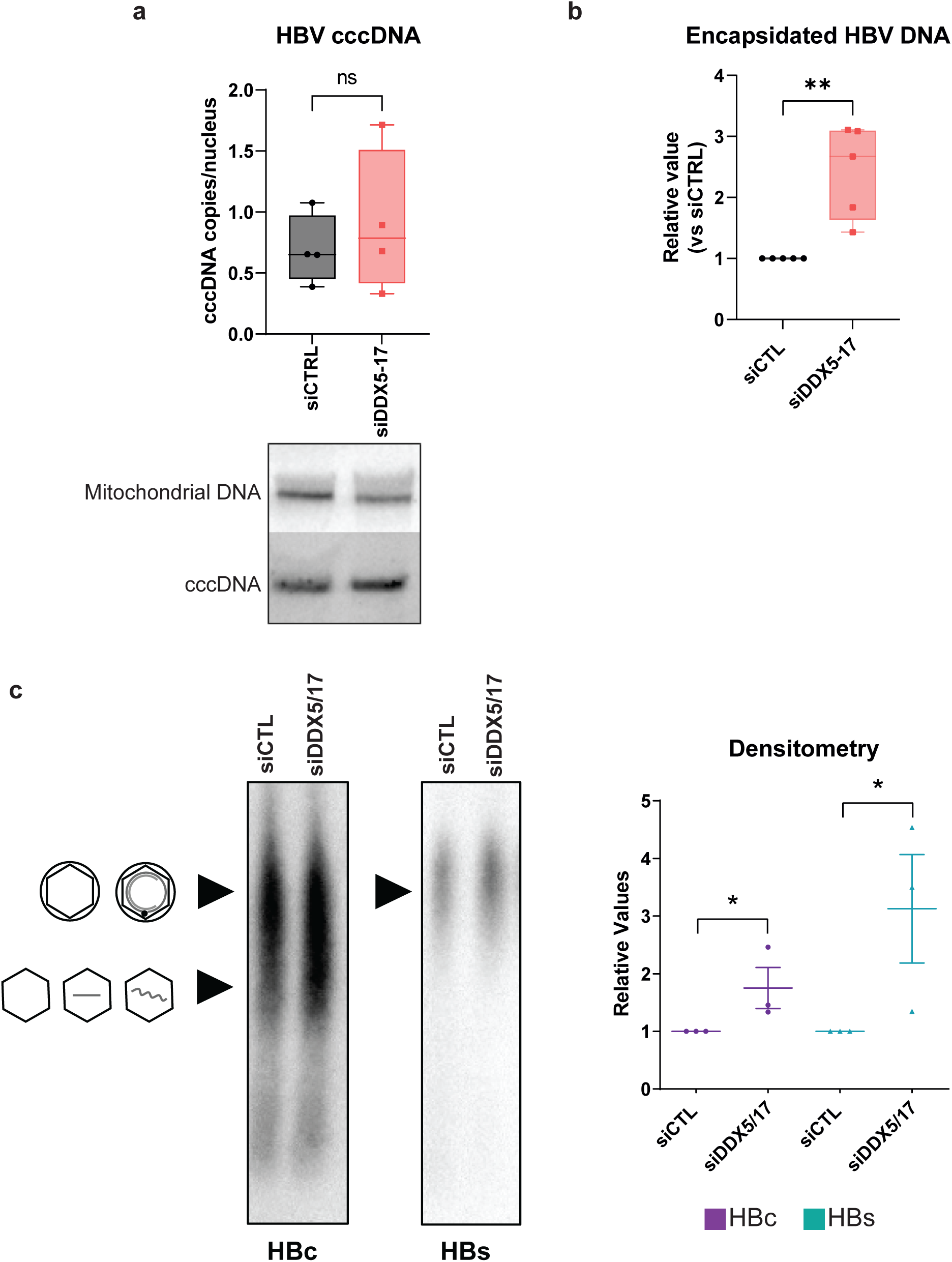
Repression of DDX5 and DDX17 increases HBV replication. HepG2-NTCP cells were infected and transfected at 4 and 6 days post-infection with control siRNA (siCTRL) or siRNAs directed against *DDX5* and *DDX17* mRNAs (siDDX5-17). Samples were collected eight days post-infection. **a** *Top panel:* DNA was isolated and digested with ExoI/ExoIII exonucleases and cccDNA was quantified by qPCR and normalized to the level of β-globin. *Bottom panel:* cccDNA and mitochondrial DNA (used as loading control) were visualized by Southern blot experiments using DIG-labelled specific probes against HBV DNAs. Uncut gel is presented in Figure S5. **b** HBV capsids obtained from the same number of cells in siCTL and siDDX5/17 were isolated and HBV DNA isolated and quantified by qPCR. The data are expressed as fold change of the siCTRL condition. **c** Native agarose gel electrophoresis (NAGE) of isolated viral particles in cell supernatant. *Left panel*: HBV particles were obtained from the same volume of lysate in each condition and detected by western-blot using an HBc-specific antibody to detect capsids (rcDNA-containing and empty virions, upper, and naked capsids, lower) and an HBsAg-specific antibody to detect enveloped particles. *Right panel*: Boxplots represent the first quartile, the median (horizontal line) and the third quartile of the densitometry analysis of 3 independent experiments and data are expressed as fold change of siCTL condition. Error bars represent the minimal and maximal values. ns: Mann-Whitney p-value > 0.05. *: p<0.05 (Kruskal-Wallis multiple comparisons test); **: Mann-Whitney p-value < 0.01.

Altogether, these data establish a functional link between DDX5 and DDX17 expression, regulation of HBV transcription termination through the recognition of cPAS sequence, HBV RNA stability and viral replication.

## DISCUSSION

Co-transcriptional 3’ end mRNA processing is critical to fine tune gene expression in eukaryotic cells. Also, the choice of PAS could play a major role in viral replication, as demonstrated for human Herpes Simplex Virus 1 (HSV-1). Indeed, alternative polyadenylation of viral transcripts contributes to an optimal viral gene expression during the lytic phase, whereas it participates to the HSV-1 immune escape during the latency stage (24). Despite the identification of a PAS in the HBV genome decades ago (4), the molecular mechanisms governing its recognition and its importance in HBV replication still remain uncovered questions.

Here, we characterized the 3’ heterogeneity of HBV transcripts at the single molecule level in *in vitro* cellular models, including primary human hepatocytes, and in liver samples derived from CHB patients. We found that the majority of HBV transcripts precisely stop 14 nucleotides downstream of the cPAS, confirming previous observations obtained from plasmid-derived HBs transcripts, which were shown to terminate between 12 and 19 nt downstream of cPAS (4). Interestingly, our results provide new information with the identification of previously overlooked HBV RNAs with a longer 3’ region, spanning around 700 bp after the cPAS, strongly suggesting the presence of a transcriptional readthrough. Using an HBx expressing vector fused to this identified 3’UTR sequence, we demonstrated that readthrough transcription was associated with shorter RNA half-life and lower HBx protein expression. Modulation of the size of 3’ UTR region is known to serve as an additional mechanism of regulation of gene expression (22). Most mechanisms proposed to explain this regulation rely on the recruitment of trans-regulators of RNA metabolism (22).

We identified DDX5 and DDX17 RNA helicases as critical promoters of HBV transcripts readthrough. Their depletion strongly decreased readthrough transcription derived from HBV natural infection and from the HBx-expressing plasmid. The lack of effect on transcription termination when cPAS was mutated, together with the ChIP and CLIP data pointing at an association of the helicases to both HBV genome and RNAs, strongly suggest that DDX5/17 act via the regulation of cPAS recognition, although we cannot completely exclude an indirect contribution of other effectors regulated by DDX5 and DDX17. The functional link we provided between DDX5/17-dependent HBV transcriptional fidelity and viral replication pointed at DDX5/17 as crucial restriction factors for HBV. The observation made by Zhang *et al.* (11) that HBV infection decreases DDX5 expression is in agreement with our data and suggests that HBV represses DDX5/17 expression to allow proper viral transcriptional fidelity and higher viral replication by decreasing the probability of having transcripts with longer 3’UTR which are less stable. It is therefore not surprising that these transcripts represent only a minority of the total HBV transcriptome in infected cells (**Figure 2**). Moreover, given the crucial role of HBx in viral replication (23), a precise termination of its transcript is thus necessary to ensure HBV infection establishment and persistence.

Both DDX5 and DDX17 were already identified as regulators of transcription termination (9, 25, 26). However, their role in HBV seems to be the opposite than on HeLa or SHSY-5Y cells. The fact that the effect of DDX5/17 on transcription termination strongly depends on the chromatin topology of the regulated locus (9) would suggest a role for HBV minichromosome topology in their effect. Unfortunately, the small size of this episome has made it challenging to unravel its tri-dimensional conformation so far.

Discordant results were obtained by different groups regarding the role of DDX5 or DDX17 in HBV replication. Indeed, DDX5 was shown to repress cccDNA transcription by stabilizing the PRC2 complex subunit SUZ12 (11). DDX17 was shown to inhibit pgRNA encapsidation by preventing the association between HBV Polymerase and the pgRNA ε-motif in HepG2 cells (12). However, Dong *et al*. recently showed opposite results, suggesting that HBx increased DDX17 expression which in turn promoted HBV transcription and replication in HepG2-NTCP cells and in a hydrodynamic HBV mouse model (13). Since DDX5 and DDX17 have redundant functions and cross-regulate themselves (8, 27), it cannot be excluded that these contrasting results might be due to an unbalanced ratio between the two helicases after downregulating only one of them. Here, the silencing of both DDX5 and DDX17 clearly demonstrated their joint role as HBV replication restriction factors.

Altogether, our data point out the pivotal importance of HBV transcription termination in HBV replication and provide a clear example of how co-transcriptional processing of HBV RNAs can modulate viral persistence. This study opens the door of a deeper understanding of the mechanisms controlling HBV transcription termination and their role in viral persistence and pathogenesis.

## Acknowledgments

We would like to thank Maud Michelet, Jennifer Molle, Anaëlle Dubois and Sarah Heintz for their help in the isolation of primary human hepatocytes, as well as Prof. M. Rivoire’s surgical staff for providing liver resections.

## Authorship contribution

Conceptualisation and formal analysis: FC, GG, BT. Funding acquisition: GG, BT, CB, FZ. Investigation: FC, GG, PH, MR, MGM, AD, JF, ML. Methodology: GG, FC. Bioinformatics: XG, HP, CC, CG and CB. Resources: CS, IC, FZ, HHV, MR. Visualisation: GG, FC, BT. Writing – original draft: GG, BT. Writing – review and editing: all authors.

