## Supplementary material and figures for "Helicases DDX5 and DDX17 promote Hepatitis B Virus transcription termination heterogeneity in infected human hepatocytes"

### EXPERIMENTAL PROCEDURES

#### Production of HBV viral inoculum

HBV (genotype D, subtype ayw) inoculum was prepared from filtered HepAD38 supernatants by polyethylene-glycol-MW-8000 (PEG8000, SIGMA) precipitation (8% final) as previously described (1). Viral stock with a titer reaching at least $1x10^{10}$ viral genome equivalents (vge)/mL was tested endotoxin free and used for infection.

#### PHH isolation

Primary human hepatocytes (PHH) were isolated from surgical liver resections after informed consent of patients (IRB agreements #DC-2008-99 and DC-2008-101) as previously described (2) and plated in complete William's supplemented with 1 % penicillin/streptomycin (Life Technologies), 1% L-Glutamine (Life Technologies), 5 µg/mL insulin (Sigma-Aldrich), 25 µg/mL hydrocortisone hemisuccinate UPJOHN (SERB) and 5 % fetal calf serum (FCS. Fetalclone II™, PERBIO). PHH were maintained in William's medium supplemented with 1.8 % DMSO (Sigma-Aldrich). All PHH-related data were obtained from at least two distinct donors. All experiments on PHH were performed as described for HepG2-NTCP cells.

#### Cell culture and HBV infection

HepG2-NTCP cells were seeded at $10^5$ cells/cm2 in DMEM medium supplemented with 1% penicillin/streptomycin (Life Technology), 1% sodium pyruvate (Life Technology), 1% glutamine (Life Technology), 5% FCS (Fetalclone II™, PERBIO). The day after, medium was renewed and complemented with 2.5% DMSO (SIGMA). After 72h, cells were infected at a multiplicity of infection of 250 in the presence of 4% PEG800 for up to 16h and then extensively washed with PBS and maintained in complete DMEM medium containing 2.5% DMSO until harvesting. Intracellular accumulation of viral RNA and DNA, were monitored by RT-qPCR, qPCR, and Southern blotting. Total DNA was purified from infected cells using MasterPure™ Complete DNA Purification Kit (Epicentre). Total RNA was extracted using ExtractAll TRI-

Reagent (MRC), precipitated in isopropanol, washed in ethanol and resuspended in RNasefree water.

**Liver samples from chronic hepatitis B patients**

Human liver samples derived from 2 untreated chronic hepatitis B (CHB) male patients belonging to a historical cohort collected at Hospices Civils de Lyon (Lyon University Hospitals, France). These patients underwent liver biopsy as part of their clinical follow-up, a fragment was preserved for research purposes and stored at -80°C. The protocol was approved by the competent Institutional Ethics Committee (CPP Sud est IV 11/040, authorization number DC-2008-235). Written informed consent was obtained from all patients and/or their legal guardians to underwent liver biopsy. No patients were co-infected with HIV, hepatitis C virus or hepatitis delta virus.

**siRNA transfection**

siRNA (siDDX5 catalog# 4392420; siDDX17 catalog# 4390824; ON-TARGETplus LUC siRNA, Dharmacon) were transfected either concomitantly (12.5 nM siDDX5, 12.5 nM siDDX17), or alone (25 nM siRNA control: siLUC) in HepG2-NTCP cells and PHH four days post-infection, using Lipofectamine RNAiMAX (ThermoFisher), following manufacturer's protocol. The day after transfection, culture medium was changed, and cells were transfected a second time six days post-infection. The culture medium was changed the day after the second transfection. The supernatants were collected and cells were harvested two days after the second transfection.

**Plasmids**

HBx fragment (1374-2625) fused to a V5 sequence at the 5' extremity was obtained by PCR using the PrimeSTAR mix 1X (Takara), 200 nM primers containing AgeI and NotI restriction sites and cDNA obtained from HBV-infected HepG2-NTCP cells (**Table S1**) following the PCR program in a CFX96 Biorad thermocycler: 3 min 95 °C, 30 cycles of 30 sec 95 °C, 30 sec 60 °C, 1 min 72 °C and a final step of 5 min 72°C. The same fragment carrying mutation of the cPAS was obtained by overlapping PCR using the same program. The PCR amplicons were then cloned between the AgeI and NotI restriction site of the pcDNA6 plasmid (gifted by M.D. Ruepp).

DDX5 and DDX17 expression plasmids were described in (3).

**Plasmid transfection**

1.5 millions of cells were plated on a 6-well plate and transfected in suspension with 2.5 µg plasmids and the TransIT-2020 transfection reagent (Mirus) according to manufacturer's instruction. Samples were collected two days post-transfection.

**qPCR and RT-qPCR**

Extracted RNA were digested with RNAse-free DNase I (Qiagen) and retro-transcribed into cDNA using SuperScript III reverse transcriptase according to manufacturer's instructions (Invitrogen, Carlsbad, USA). cccDNA was quantify after ExoI + ExoIII endonuclease (Epicentre) digestion of total extracted DNA for 2 hours at 37°C, followed by 20 minutes inactivation at 80°C. Real-time qPCR for total HBV RNA/DNA and cccDNA was performed using an Applied QuantStudio 7 machine (BioSystem) and TaqMan Advanced Fast Master Mix or SYBR Green Master Mix (see primers and probes below, table 1). Serial dilutions of a plasmid containing an HBV monomer (pHBV-EcoR1) served as quantification standard for HBV DNA and cccDNA. The number of cellular genomes was determined by using the β-globin gene kit (Roche Diagnostics, Manheim, Germany). RT-qPCR were analyzed using the ΔΔCt method were ΔCt = Ct (target) − Ct (GUS), where GUS is the housekeeping gene.

**Western-Blot**

Cells were lysed in RIPA buffer supplemented with 1X PIC and 1X PMSF. 20 µg of protein were migrated in 4-20% mini-PROTEAN@ TGX stain-FreeTM Precast Gel (Bio-Rad Laboratories) and transferred onto a nitrocellulose membrane (Bio-Rad Laboratories). Membranes were blocked 1 hour with 5% milk in TBS (1 x Tris Buffer Saline (Sigma)) and stained with primary antibodies in blocking buffer overnight at 4 °C (see Table XX). After primary antibody incubation, membranes were washed 3 times with TBS-T 0.1% (1X TBS with 0.1% Tween 20), stained with HRP-conjugated secondary antibodies (1/5000) for 1 hour at room temperature and washed again 3 times with TBS-T 0.1%. The detection was done using Clarity Western ECL and the ChemiDoc XRS system (Biorad).

**Nascent RNA Capture**

Cultured cells were pulsed for 2 hours with 0.2 mM Ethynil-Uridine. Total RNA were extracted as above. 1 µg of RNA was used to perform nascent RNA capture using Click-iT® Nascent RNA Capture Kit (ThermoFisher), according to the manufacturer's instructions. Precipitated RNA was retro-transcribed using Superscript IV Vilo (ThermoFisher), and qPCR was performed as described above.

**3'RACE-PCR**

RNA were extracted as described above. 1 µg of extracted RNA was digested with RNAse-free DNase I (Qiagen), purified with acid phenol (pH 4.3), precipitated with 100% ethanol, 10% Sodium Acetate (Sigma), and 1 µL Glycogen, overnight at -20°C, washed with 70% ethanol and resuspended in water. A 3'-DNA-Adaptor Ligation was performed with T4 RNA ligase 2 truncated (NEB) in presence of RNase-OUT (ThermoFisher) for 1 hour at 25°C. After acid phenol precipitation, gene specific reverse-transcription of anchored RNA was performed using ProtoScript II (NEB). The final PCR reaction was done with Platinum SuperFi II DNA Polymerase (ThermoFisher) (see primers Table 1). PCR products were deposited and migrated on a 1% ultrapure agarose gel.

**Nanopore sequencing**

3'RACE-PCR amplicons were further processed with the PCR barcoding kit (SQK-PBK004, Oxford Nanopore Technologies) according to the manufacturer's instruction. Sequencing was performed on a MinION Single FlowCell (FLO-MIN106D, Oxford Nanopore Technologies) for 48 h. Analysis was performed as described in (4). *De novo* assembly was performed to generate a consensus sequence, which was blasted (https://blast.ncbi.nlm.nih.gov/Blast.cgi) to check for homology with HBV genome. This consensus sequence was used as a reference genome to align the Nanopore reads. Integrated Genomics Viewer (IGV) was used to visualized the alignment files (5, 6).

**Cross-Link Immunoprecipitation (CLIP)**

CLIP experiments were carried out 7 days post-infection. Cells were washed in 1X PBS and UV cross-linked at 200 mJ/cm2. Total cell lysate was obtained by 20 min RIPA lysis buffer incubation on ice, followed by sonication. Total cell lysate was pre-cleaned in RIPA buffer with Protein G-coupled magnetic beads (Dynabeads, Invitrogen), and then subjected to overnight immunoprecipitation at 4 °C using 2–4 µg of antibodies (see Table 2). Chromatin incubated with no antibody or anti-H3.3 antibody was processed as other conditions and was used as negative controls. Immune complexes were then incubated 2 hours with Protein G-coupled magnetic beads at 4 °C, washed, and eluted in ChIP elution buffer. The flow-through from the no antibody condition was used as input. Immunoprecipitated RNA were extracted using ExtractAll TRI-Reagent (MRC), precipitated in isopropanol, washed in ethanol and resuspended in RNase-free water. RT-qPCR was performed as described above. Samples were normalized to input RNA using the ΔΔCt method were ΔCt = Ct (input) – Ct (immunoprecipitation) and calculated as percentage of the input.

**Chromatin Immunoprecipitation**

ChIP experiments were carried out 7 days post-infection. Cells were washed in 1X phosphate buffered saline (PBS), and incubated for 15 minutes with 1% formaldehyde at 37 °C and quenched with 0.125 M Glycine for 10 minutes at 37°C. For nuclear extracts preparation, cells were lysed in lysis buffer (5 mM PIPES, 85 mM KCl, 0,5% NP-40, 1 mM PMSF, 1X Protease inhibitor cocktail (PIC) (ThermoFisher)). After “tight” douncing (10 times) and 5 minutes centrifugation at 3000 rpm, nuclei were resuspended in sonication buffer (1% SDS, 10 mM EDTA, 50 mM Tris-HCl pH8, 1 mM PMSF, 1X PIC). After sonication, chromatin was pre-cleaned in RIPA buffer with Protein G-coupled magnetic beads (Dynabeads, Invitrogen), and then subjected to overnight immunoprecipitation at 4 °C using 2–4 μg of antibodies (see Table 2). Chromatin incubated with no antibody or anti-E2F antibody was processed as other conditions and was used as negative controls. Immune complexes were then incubated 2 hours with Protein G-coupled magnetic beads at 4 °C, washed, and eluted in 10 mM Tris-HCl pH8, 5 mM EDTA, 50 mM NaCl, 1% SDS, 50 μg Proteinase K, 1X PIC. The flow-through from the no antibody condition was used as input. Immunoprecipitated DNA was extracted with phenol-chloroform isoamyl (25:24:1) and quantified by qPCR using cccDNA specific primers (see table 1). Samples were normalized to input DNA using the ΔΔCt method were ΔCt = Ct (input) – Ct (immunoprecipitation) and calculated as percentage of input.

**RNA-Seq**

RNA was extracted as above. Extracted RNAs were digested with RNAse-free DNase I (Qiagen) and tested for RNA quality control using a Bioanalyzer Instrument (Agilent). mRNA libraries were prepared using the TruSeq Stranded Total RNA Library Prep Gold (Illumina) kit to increase the depth of sequencing. Each library was paired-end sequenced to an average of 50 million reads per sample using 75x2 cycles on a NextSeq 500 Illumina platform. On average, approximately 90 million reads were generated per sample. Sequencing reads were mapped to the H.sapiens reference genome (hg19). Gene expression analyses were carried out as described in (7). Data were deposited on NCBI-GEO repository with the identifier number GSE239571.

**SUPPLEMENTARY FIGURE LEGENDS**

**Figure S1 (related to Figure 1)**

**a** Agarose gel electrophoresis profile of amplicons obtained from 3' RACE-PCR experiments performed with (RT+) or without (RT-) reverse transcriptase or with the H2O negative control. A 1 kb+ DNA ladder (L) was migrated in parallel. The figure is representative of three (HepG2-NTCP) and two (PHH) independent replicates, respectively.

**b** Graphical representation of the length of the reads obtained from the MinION single molecule sequencing of the 3'RACE amplicons of the HBV-infected HepG2-NTCP (green dots) and HBV-infected PHH (yellow dots)

**c** Blast of the de novo assembly generated consensus sequence used as a reference genome for the alignment of the single molecule sequencing.

**d** IGV view of the Nanopore sequencing reads aligned to the *de novo* assembly generated HBV consensus reference genome and obtained from HepG2-NTCP cells (green) and PHH cells (yellow) infected with HBV for 8 days. These views are from an independent biological replicate and from a different donor compared to Figure 1c. The scale is a logarithmic scale.

**e** Table showing the position and the sequence of potential PAS based on the consensus sequence of the 12 mostly used PAS in human genes (Beaudoing, 2000). The letters in bold are those always observed in these 12 PAS. The 2 PAS in yellow have been already identified and corresponds to the canonical PAS (cPAS) and to the PAS used from HBV integrated genomes (aPAS1). The PAS in blue is located 23 nucleotides upstream of the major end of the readthrough transcripts.

**f** Agarose gel electrophoresis profile of amplicons obtained from 3' RACE-PCR experiments performed with (RT+) or without (RT-) reverse transcriptase or with the H2O negative control. A 1 kb+ DNA ladder (L) was migrated in parallel.

**Figure S2 (related to Figure 2)**

**a-e** HepG2-NTCP cells were infected with HBV for 8 days and transfected at 4 and 6 days post-infection with either control siRNA (siCTL) or siRNAs directed against *DDX5* and *DDX17* mRNA (siDDX5-17).

**a** Proteins were extracted and Western blot experiments were conducted using an anti-DDX5, an-anti-DDX17 (top panels) or an anti-ß-actin (ACTB, bottom panel, loading control). The figure is representative of 10 independent biological replicates.

**b** RT-qPCR quantification of the *DDX5* and *DDX17* mRNAs levels. The signals obtained for DDX5 and DDX17 were normalized to the level of the *GUS* mRNAs and to the signals of siCTL condition. The bars represent the average of 10 independent biological replicates. The error bars represent the standard error of the mean. ****: p-value < 0.001 from a Mann-Whitney test comparing siCTL and siDDX5-17 conditions.

**c** Electrophoresis profile of amplicons obtained from 3' RACE-PCR experiments performed from HBV RNAs isolated from HBV-infected HepG2-NTCP cells transfected with either a control siRNA or siRNAs directed against *DDX5* and *DDX17* mRNAs. A 100 bp DNA ladder was run in parallel.

**d** Graphical representation of the length of the reads obtained from the MinION single molecule sequencing of three independent biological replicate of the 3'RACE amplicons.

**e** Frequency of the reads longer (in red) or shorter (in blue) than 500 bp obtained from the single molecule sequencing of the 3'RACE amplicons mentioned in **d**.

**f** IGV view of the Nanopore sequencing reads aligned to the *de novo* assembly generated HBV consensus reference genome. The figure represents two other independent biological replicates compared to Figure 2f. The scale is a logarithmic scale.

**g-k** PHH were infected with HBV for 8 days and transfected at 4 and 6 days post-infection with either a control siRNA (siCTL) or siRNAs directed against *DDX5* and *DDX17* mRNA (siDDX5-17).

**g** Proteins were extracted and Western blot experiments were conducted using an anti-DDX5, an-anti-DDX17 (top panels) or an anti-ß-actin (ACTB, bottom panel, loading control). The figure is representative of 6 independent biological replicates.

**h** RT-qPCR quantification of the *DDX5* and *DDX17* mRNAs levels. The signals obtained for DDX5 and DDX17 were normalized to the level of the *GUS* mRNAs and to the signals of siCTL condition. The bars represent the average of 6 independent biological replicates. The error bars represent the standard error of the mean. *: p-value < 0.05 from a Mann-Whitney test comparing siCTL and siDDX5-17 conditions.

**i** Electrophoresis profile of amplicons obtained from 3' RACE-PCR experiments performed from HBV RNAs isolated from HBV-infected PHH cells transfected with either a control siRNA or siRNAs directed against *DDX5* and *DDX17* mRNAs. A 100 bp DNA ladder was run in parallel.

**j** Graphical representation of the length of the reads obtained from the MinION single molecule sequencing of the 3'RACE amplicons. These data are from a second PHH donor compared to Figure 2.

**k** Barplots indicating the frequency of the reads longer (in red) or shorter (in blue) than 500 bp obtained from the single molecule sequencing of the 3'RACE amplicons mentioned in **j**.

**l** IGV view of the Nanopore sequencing reads aligned to the *de novo* assembly generated HBV consensus reference genome. The figure represents two other independent biological replicates compared to Figure 2f. The scale is a logarithmic scale.

**Figure S3 (related to Figure 4)**

HepG2-NTCP cells were transfected with pcDNA6 plasmids described in **Figure 4a**. Two days after, DNA was extracted and qPCR experiments were performed to quantify the pcDNA6 plasmid whose signal has been normalized to the host locus *SLC10A1*. Each dot represents an individual biological replicate. Boxplots represent the first quartile, the median (horizontal line) and the third quartile of six independent biological replicates. Error bars represent the minimal and maximal values. ns: Mann-Whitney p-value > 0.05.

**Figure S4 (related to Figure 5)**

**a** HepG2-NTCP cells were transfected with control siRNA (siCTRL) or siRNAs directed against *DDX5* and *DDX17* mRNAs one day prior to their transfection with the pcDNA6 plasmids expressing HBx transcripts carrying (cPASmut) or not (cPASwt) mutations in the cPAS. Two days after, proteins were extracted and subjected to Western blot analyses using either an anti-DDX17 (top blot) or an anti-DDX5 (middle blot) antibody. Total proteins (bottom blot) were revealed and used as loading controls. The blots are representative of four independent biological replicates.

**b-e** HepG2-NTCP cells were transfected with plasmids encoding HA-DDX5 or HA-DDX17 either alone or in combination (HA-DDX5/17) or GFP one day prior to their transfection with the pcDNA6 plasmids expressing HBx transcripts carrying (cPASmut) or not (cPASwt) mutations in the cPAS. Two days after, proteins and RNAs were extracted.

**b** Proteins were subjected to Western blot analyses using either an anti-DDX17 (top blot) or an anti-DDX5 (middle blot) antibody (**d**). Total proteins (bottom blot) were revealed and used as loading controls. The blots are representative of five independent biological replicates.

**c, e** RNAs were subjected to RT-qPCR analyses to quantify long (**c**) and total (**e**) HBx transcripts. Long HBx transcripts levels were normalized to the signal obtained for total HBx transcripts while total HBx transcripts levels were normalized to *GAPDH* host mRNA levels. Both were normalized to the signal obtained in the cPASwt-GFP condition

**d** *Top panel:* Western blot analyses using an anti-V5 antibody (top blot). Total proteins (bottom blot) were revealed and used as loading controls. The blots are representative of five independent biological replicates. *Bottom panel:* V5 (HBx) signal was quantified and normalized to the corresponding total proteins load and GFP-cPASwt conditions.

**c-e** Each dot represents one replicate. The blox plot represent the first quartile, the median (horizontal line) and the third quartile of five independent biological replicates. Error bars represent the minimal and maximal values. ns: Mann-Whitney p-value > 0.05; *: Mann-Whitney p-value < 0.05. **: Mann-Whitney p-value < 0.01.

**Figure S5 (related to Figure 7)**

**a-b** Full gel image of the Southern blot displayed in **Figure 7a** obtained after incubation of the membrane with HBV-specific probes (**a**) or with mitochondrial ND2 probes (**b**)

a

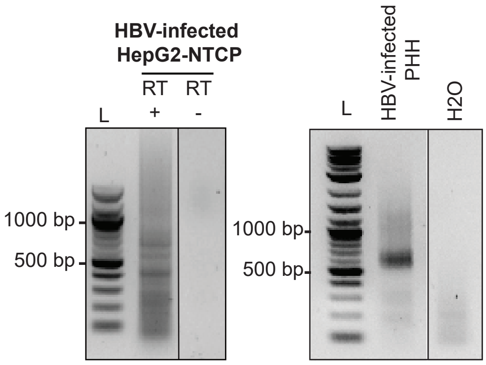


b

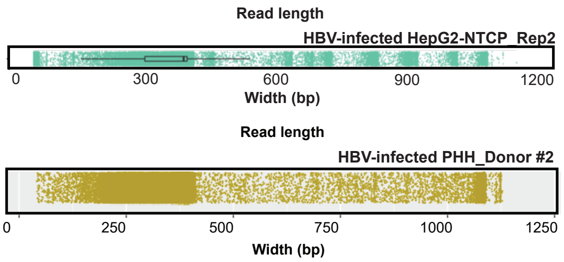


c

**Hepatitis B virus complete genome, genotype D**

Sequence ID: AJ344117.1 Length: 3182 Number of Matches: 1

Range 1: 1562 to 2631 GenBank Graphics

| Score | Expect | Identities | Gaps |
|---|---|---|---|
| 1877 bits(1016) | 0.0 | 1053/1070(98%) | 6/1070(0%) |

d

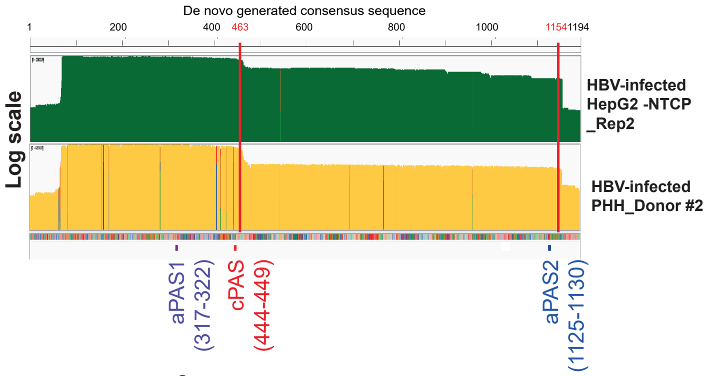


e

| Consensus Reference Genome Coordinates | | HBV Genome Coordinates | | | |
|---|---|---|---|---|---|
| Start | End | Start | End | Sequence | Comments |
| 179 | 684 | 1651 | 1656 | CTTACA | |
| 252 | 257 | 1724 | 1729 | TTTAAA | |
| 281 | 286 | 1754 | 1759 | ATTAGA | |
| 286 | 291 | 1759 | 1764 | ATTAAA | |
| 317 | 322 | 1790 | 1795 | CATAAA | aPAS1 = HBV integrated genome PAS |
| 444 | 449 | 1918 | 1923 | TATAAA | cPAS |
| 518 | 523 | 1992 | 1997 | TCTAGA | |
| 551 | 556 | 2025 | 2030 | CTTAGA | |
| 667 | 672 | 2143 | 2148 | TCTAGA | |
| 708 | 713 | 2184 | 2189 | CCTAAA | |
| 769 | 774 | 2245 | 2250 | GTTATA | |
| 771 | 776 | 2247 | 2252 | TATAGA | |
| 820 | 825 | 2296 | 2301 | TATAGA | |
| 891 | 896 | 2368 | 2373 | CCTAGA | |
| 1072 | 1077 | 2549 | 2554 | AATATA | |
| 1074 | 1079 | 2551 | 2556 | TATACA | |
| 1080 | 1085 | 2557 | 2562 | TTTACA | |
| 1125 | 1130 | 2603 | 2608 | CTTACA | aPAS2 |

f

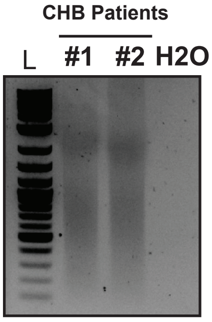


**Figure S1 (related to Figure 1)**

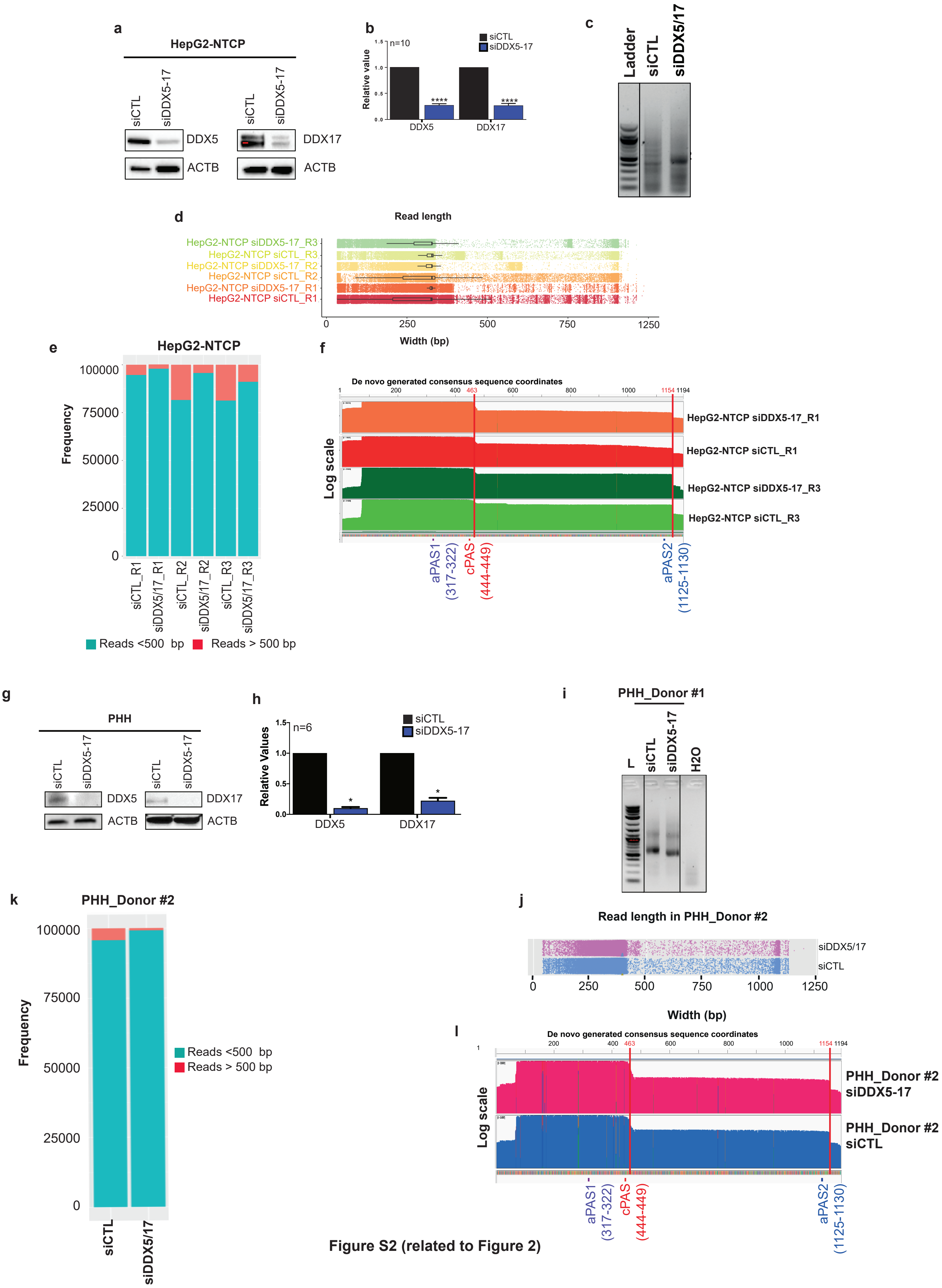


**Figure S2 (related to Figure 2)**

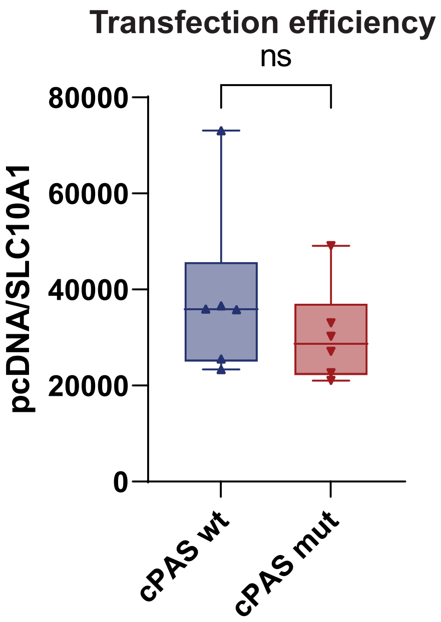


Figure S3 (related to Figure 4)

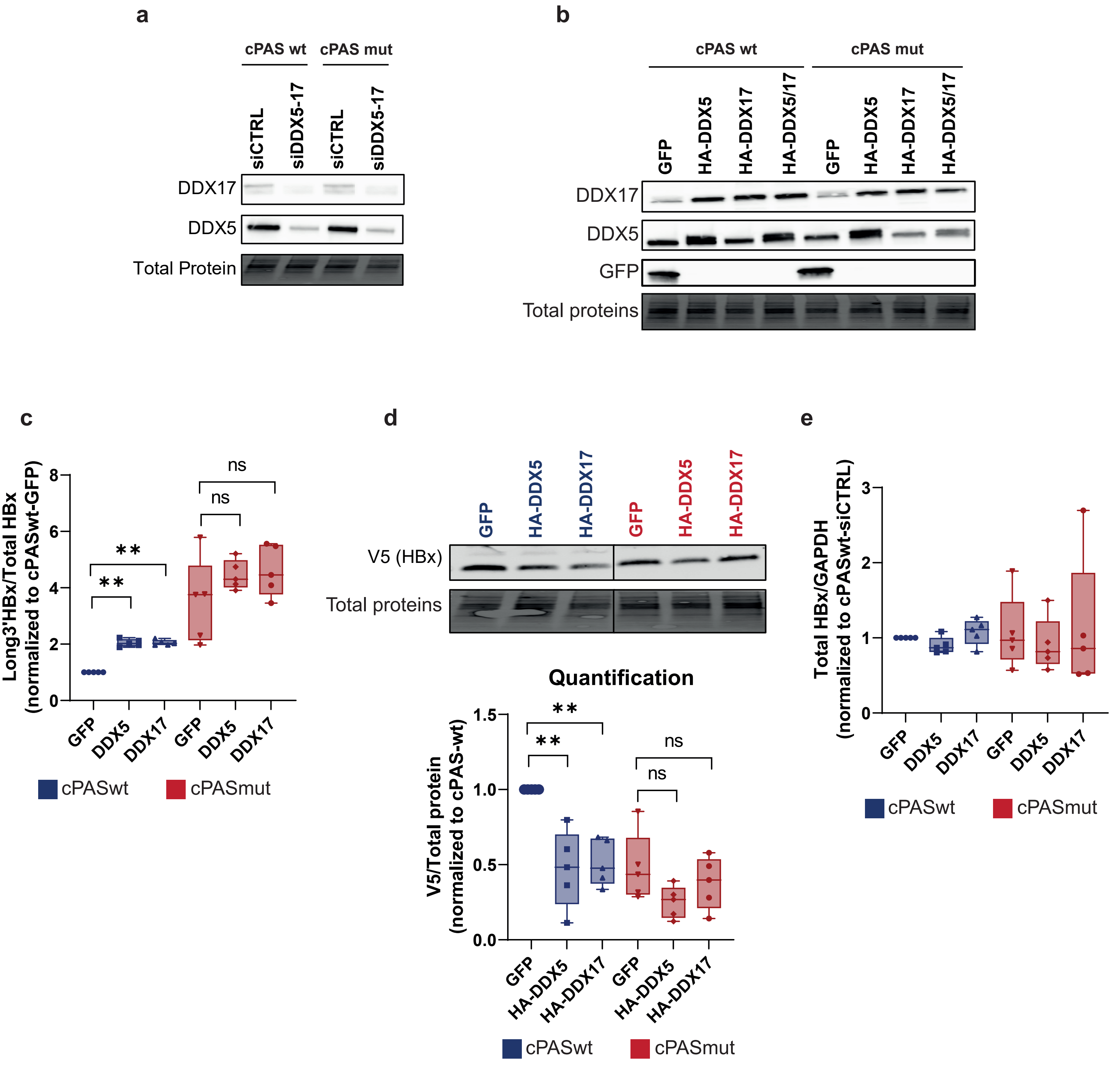


**Figure S4 (related to Figure 5)**

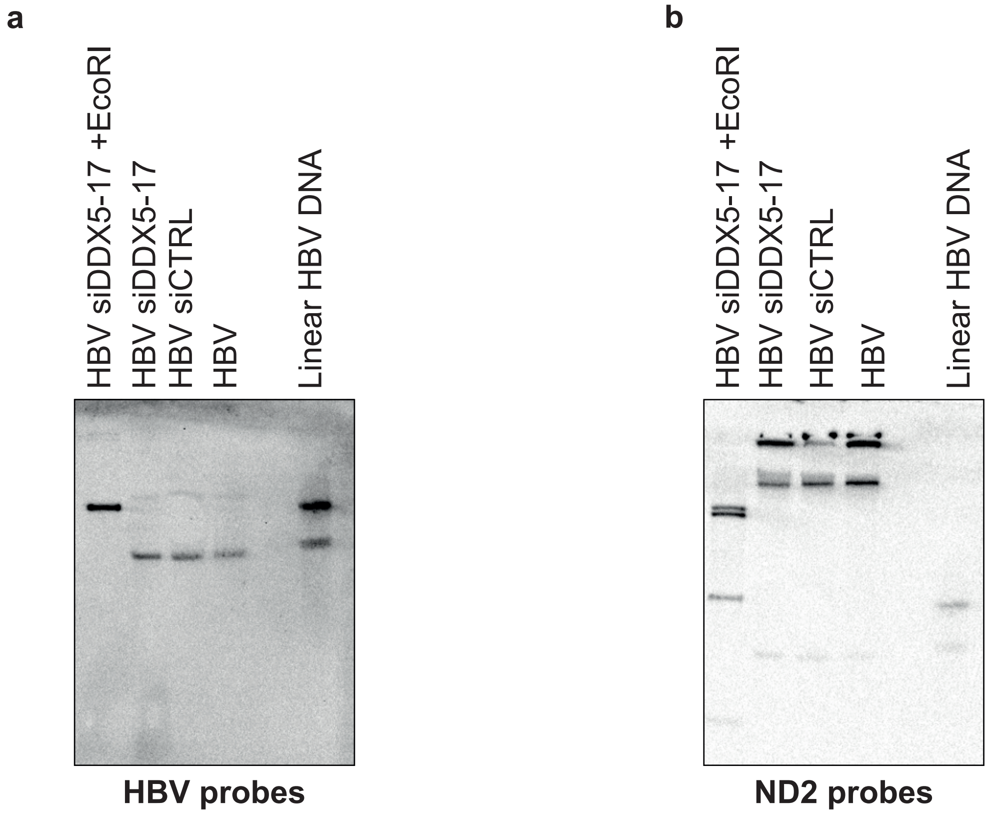


**Figure S5 (related to Figure 7)**
